# GenMasterTable: A user-friendly desktop application for filtering, summarising, and visualising large-scale annotated genetic variants

**DOI:** 10.1101/2025.04.10.648172

**Authors:** Jing Zhai, Nelly Pitteloud, Federico A. Santoni

**Author notes:** Federico A. Santoni and Jing Zhai are co-corresponding authors.

## Abstract

**Background:** The rapid expansion of next-generation sequencing (NGS) technologies has generated vast amounts of genomic data, creating a growing demand for secure, scalable, and accessible tools to support variant interpretation. However, many existing solutions are command-line based, rely on cloud or server infrastructures that may pose data privacy risks, lack flexibility in supporting both VCF and CSV formats, or struggle to handle the scale and complexity of modern genomic datasets. There is a clear need for a user-friendly, locally operated application capable of efficiently processing annotated variant data for large-scale cohort level analysis.

**Results:** We introduce GenMasterTable, a free, secure, and cross-platform desktop application designed to simplify variant analysis through an intuitive graphical user interface (GUI). As the first tool to enable comprehensive cohort-level analysis from both VCF and CSV files, GenMasterTable provides advanced functionality for merging, filtering, summarizing, and visualizing large-scale annotated datasets. Tailored for users without programming expertise, it enables rapid and accurate exploration of genetic variants, making it a practical solution for both research and clinical settings.

**Conclusion:** GenMasterTable addresses critical limitations in current variant analysis workflows by combining usability, data security, and scalability. Its support for multiple input formats and locally executed operations empowers clinicians, geneticists, and researchers to perform comprehensive variant analysis efficiently without the need for programming expertise.

## Background

The advent of next-generation sequencing (NGS) technologies has led to an unprecedented surge in genomic data. Analysing these large datasets is crucial for identifying pathogenic variants, understanding disease mechanisms, and translating genomic findings into clinical practice. Genomic datasets, such as whole-genome sequencing (WGS), whole-exome sequencing (WES), and RNA sequencing (RNA-seq), often derived from patient samples, are not only massive in scale but also highly sensitive, necessitating robust security measures to ensure data privacy and compliance with regulatory standards.

Standard bioinformatics pipelines, such as those based on the Genome Analysis Toolkit (GATK), followed by annotation tools like Variant Effect Predictor (VEP) (McLaren et al., 2016), ANNOVAR (Wang et al., 2010), or FAVOR (Zhou et al., 2023), typically produce results in Variant Calling Format (VCF) files or structured tabular formats (e.g., CSV/TSV). While VCF files have long been the standard, modern annotation tools increasingly favor tabular formats due to their flexibility. However, both formats present significant challenges in terms of data management and analysis.

Existing tools for handling VCF files, such as GEMINI (Paila et al., 2013), Variant tools (Wang et al., 2014), SnpSift (Cingolani et al., 2012b) often require command-line expertise or programming skills, limiting their accessibility to clinicians and geneticists. Moreover, most of these tools are web-based, requiring users to upload sensitive genomic data to external servers such as myVCF (Pietrelli et al., 2017), BrowseVCF (Salatino et al., 2017), VCF-Miner (Hart et al., 2016), VCF-Explorer (Akgun et al., 2017), VCF-Server (Jiang et al., 2019), raise concerns about privacy and regulatory compliance under certain conditions. A notable exception is Cutevariant (Schutz et al., 2021), a standalone desktop application that offers a graphical user interface (GUI) and ensures local data processing. However, Cutevariant relies on a specialized query language (VQL), which can be a hurdle for non-technical users who are unfamiliar with SQL-like syntax. Additionally, it supports only VCF files as input, limiting its flexibility.

For tabular data (CSV), standard database solutions such as SQLite(https://www.sqlite.org), Oracle(https://www.oracle.com), PostgreSQL(https://www.postgresql.org), Spark (Zaharia et al., 2010), and Google BigQuery (https://cloud.google.com/bigquery) might provide extensive genomic data management capabilities. However, these solutions require proficiency in SQL and other technical skills, making them impractical for many researchers and clinicians. On the other hand, widely used desktop tools like Microsoft Excel (https://www.microsoft.com/en-us/microsoft-365/excel), provide more intuitive interfaces to process tabular structured data but fall short due to inherent limitations in handling large-scale genomic datasets, leading to performance bottlenecks and restricted scalability.

To address these challenges, we present GenMasterTable, a user-friendly, desktop-based application compatible with most diffuse operating systems. It features an intuitive graphical user interface (GUI) designed to facilitate the merging, advanced filtering, summarization, and visualization of large-scale annotated genetic variant data at the cohort level. GenMasterTable supports multiple input formats, including both VCF and CSV files derived from DNA or RNA sequencing. Users can upload individual files or merge multiple files into a single dataset, enabling cohort-level analysis even when sequencing data are provided on a per-sample basis.

Unlike many existing solutions, GenMasterTable operates entirely locally, ensuring robust data security and compliance with regulatory requirements—critical for handling sensitive patient information. It requires no programming or database expertise, making comprehensive variant analysis accessible to clinicians, geneticists, and researchers across disciplines.

In comparison with the previously mentioned existing solutions, GenMasterTable supports both VCF and CSV formats while enabling comprehensive cohort-level variant analysis through an intuitive graphical interface—without requiring any programming skills. It bridges the gap between high-throughput bioinformatics pipelines and user-friendly tools tailored for clinical and translational research.

## Methods

GenMasterTable is a standalone, desktop-based application developed in Python for efficient genomic data processing. The backend integrates packages such as Pandas for high-performance data manipulation, NumPy for numerical computations, PyVCF for VCF file parsing, and the re module for advanced text-based filtering. The graphical user interface (GUI) is implemented using Tkinter, incorporating an embedded Panda stable-based table viewer to facilitate interactive data exploration.

The application follows a multi-panel PanedWindow layout, optimizing the user experience by enabling seamless interaction between data visualization and control functionalities. The table frame displays datasets in an interactive spreadsheet format, while the control frame provides filtering options. Users can import and merge multiple CSV and VCF files, apply column-based filters, and dynamically update data views.

GenMasterTable is deployed as a standalone executable (.app for macOS, .exe for Windows) using PyInstaller, ensuring ease of installation and use without requiring prior software dependencies.

### Data Processing and Parsing

GenMasterTable supports the loading of individual files and the merging of multiple annotated VCF or CSV files into a single dataset, facilitating the analysis of next-generation sequencing (NGS) data at a cohort level, where annotated files are generated on a per-individual basis (patient or control) using a standard bioinformatics pipeline.

To facilitate the handling of large-scale genomic data, GenMasterTable employs batch processing techniques, including VCF parsing through PyVCF with chunk-based extraction of key attributes such as chromosome position, alleles, genotype calls, quality scores, and INFO fields. Similarly, CSV files are processed in a chunked reading approach to mitigate memory overhead, ensuring efficient performance on extensive datasets. All data is structured within a Pandas DataFrame, enabling high-performance filtering, merging, and visualization.

### Graphical User Interface and Filtering System

The GUI is structured around a multi-panel layout, featuring an interactive variant table display, a file operations panel, and a filtering control section. Users can select columns and define filtering criteria through a drop-down menu and text fields, enabling multi-tier filtering. The filtering engine is optimized using vectorized Pandas functions, allowing for rapid execution of both single-column queries and complex multi-column filters on datasets containing millions of variants. For categorical data, filtering is designed to support lists of values, while a toolbar on the right side of the GUI provides an additional filtering option for column-specific queries. This feature is particularly useful for defining threshold-based filters (e.g., > or <) for quantitative data, such as pathogenicity scores. Figure 1 presents a demonstration of GenMasterTable’s filtering system, applied to artificially generated test dataset.

**Figure 1:**
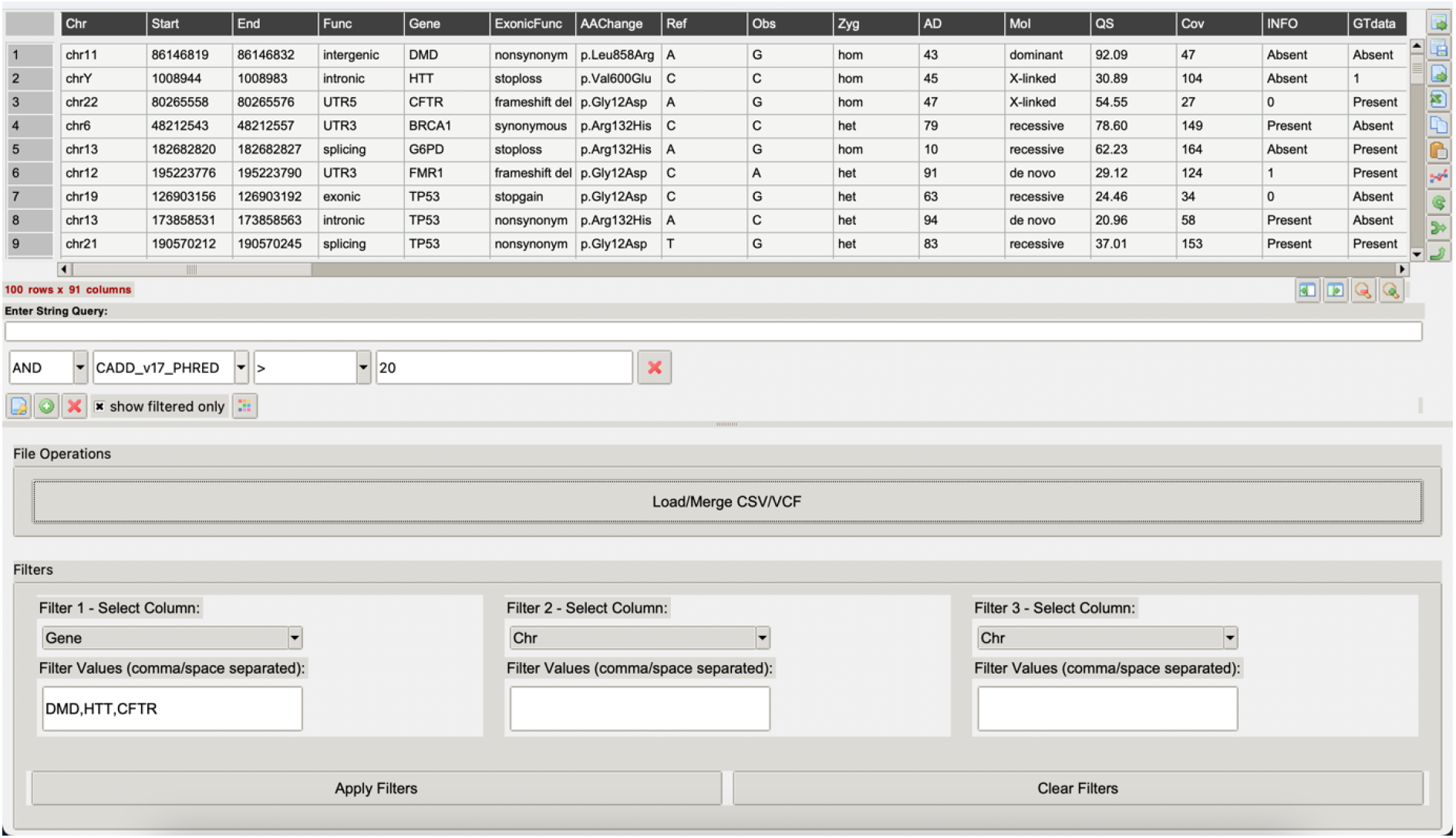
GenMasterTable’s filtering functionality using artificially generated test data.

### Interactive Data Processing

GenMasterTable offers a suite of interactive data manipulation functions designed to enhance the exploration and analysis of genomic variant datasets. Users can sort columns (e.g., genes, chromosomal positions, allelic depth) to organize data systematically, while the indexing function allows setting key identifiers, such as genes or genomic coordinates, as primary indices for streamlined analysis.

The software provides various data transformation tools, including options to delete columns and fill missing data. Value counts generate frequency distributions of variants across genes, subject IDs or other functional annotations, aiding in the identification of recurrent variants. The set data type function allows conversion between numeric, categorical, and textual formats, ensuring compatibility during the filtering process. Users can also refine dataset structures through formatting options, such as adjusting decimal precision and modifying column organization for better visualization. For detailed procedural guidance and best practices, refer to the handbook on GenMasterTable GitHub, which provides comprehensive instructions and a demo using synthetically generated genomic data to illustrate the GenMasterTable’s functionalities and workflow.

### Data Export and System Compatibility

GenMasterTable allows filtered subsets, as well as merged CSV and VCF files, to be copied and pasted directly into an open Excel sheet, preserving all annotations for downstream analysis. It is fully compatible with both Windows and macOS, ensuring broad accessibility for researchers and clinicians.

### Data for Demonstrating the Application

To demonstrate the functionality and versatility of GenMasterTable, we utilized synthetic genomic variant datasets in both VCF and CSV formats, generated using a custom Python script. These simulated datasets include a wide range of annotations commonly used in variant interpretation, such as allele frequency, predicted consequence, clinical significance, and supporting literature references. Designed to emulate real-world sequencing data, the synthetic files are publicly available in the GitHub repository to support testing, demonstration, and reproducibility.

## Results

### Benchmark

To evaluate the performance of GenMasterTable in handling large-scale variant datasets, we conducted a benchmark comparison against two alternative desktop tools: CuteVariant and Microsoft Excel. Both tools are more limited in flexibility and scope compared to GenMasterTable. While CuteVariant only supports VCF input, GenMasterTable accommodates both VCF and CSV formats, enabling broader applicability across diverse workflows. To ensure a fair comparison, all tools were benchmarked using VCF files.

As illustrated in Figure 2, GenMasterTable demonstrated significantly faster total execution times across all tested dataset sizes, ranging from 10^4^ to 10^8^ variants. For datasets exceeding 10^6^ variants, GenMasterTable remained fully responsive, while Excel reached its practical data limit and CuteVariant exhibited a sharp increase in runtime. Once data were fully loaded, GenMasterTable showed a clear advantage in loading efficiency, significantly reducing the overall time required to begin analysis.

**Figure 2:**
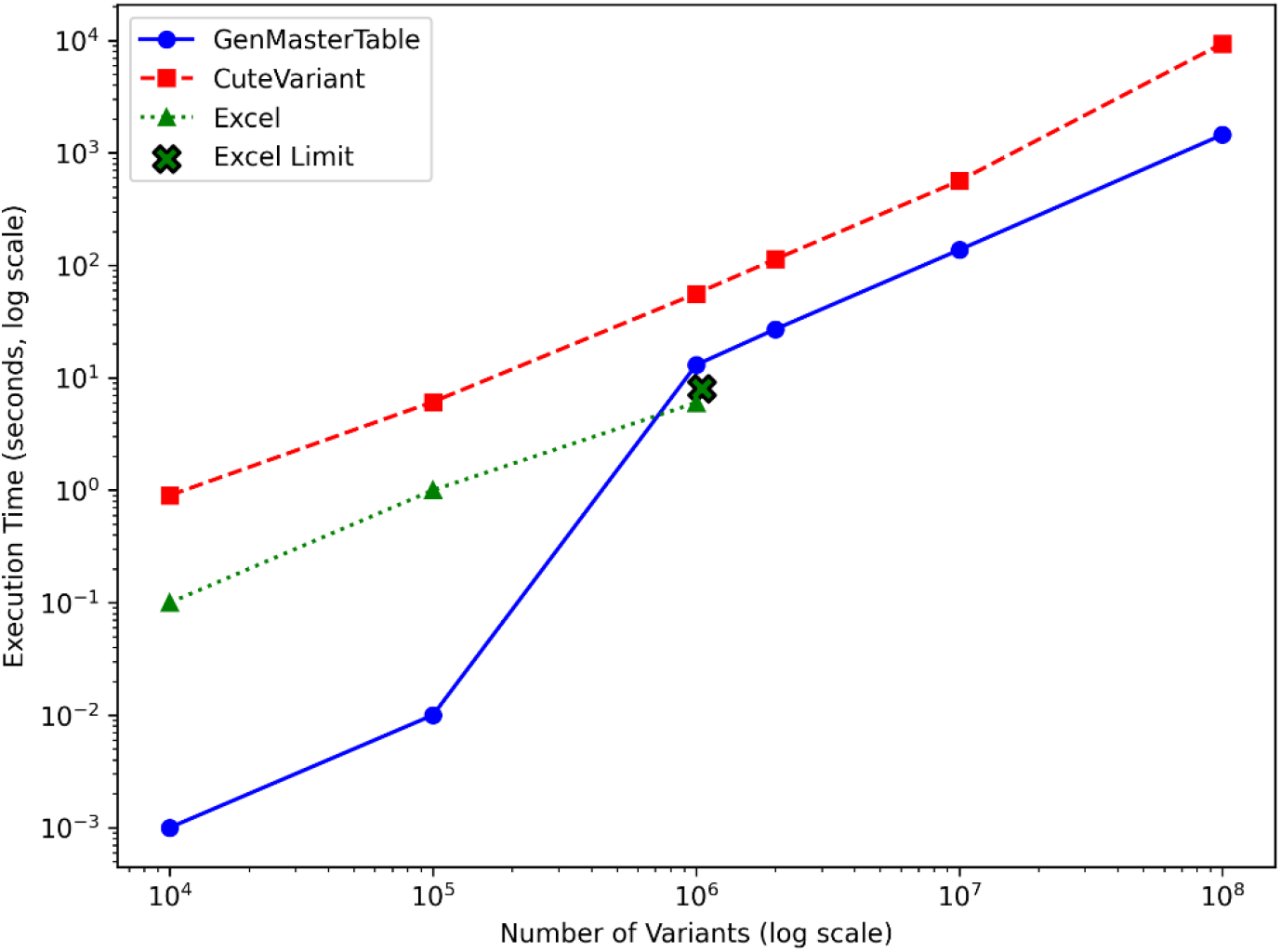
Comparative analysis of data loading times for GenMasterTable, CuteVariant, and Excel across escalating variant dataset sizes.

We further evaluated query execution performance using a synthetic whole-genome sequencing VCF file, artificially generated to resemble typical data from a healthy individual. This file contained approximately 4 million variants. When applying a basic filter such as QUAL > 30 or DP > 30, GenMasterTable returned results within 3 seconds, whereas Cutevariant required 30 seconds to complete the same operation.

All benchmarks were conducted on a Windows 10 Enterprise (64-bit) system with an Intel Core i5-11500 processor and 80 GB RAM. These results underscore GenMasterTable’s superior end-to-end efficiency, scalability, and responsiveness, making it a robust and practical solution for high-throughput genomic data analysis.

## Conclusions

GenMasterTable offers a powerful, scalable, and user-friendly solution for analysing large-scale annotated genetic variants from both individual and merged VCF/CSV files. By eliminating the need for programming expertise, ensuring local data security, and providing efficient filtering and summarization capabilities, it addresses a critical gap in genomic data analysis tools. Additionally, GenMasterTable is completely free for research use, making it an accessible alternative to commercial solutions. Its positive reception among researchers and clinicians underscores its potential as a valuable resource in genomic research and precision medicine.

## Availability and requirements

Project name: GenMasterTable.

Project home page: https://github.com/strawberrybeijing/GenMasterTable.

Operating system(s): Windows or MacOS.

Programming language: Python. Other requirements: non.

License: MIT.

Any restrictions to use by non-academics: licence needed.

## Declarations

### Abbreviations

*NGS*: Next-generation sequencing
*WES*: Whole-exome sequencing
*WGS*: Whole-genome sequencing
*RNA-seq*: RNA sequencing
*GATK*: Genome Analysis Toolkit
*VEP*: Variant Effect Predictor
*FAVOR*: Functional Annotation of Variants Online Resource
*VCF*: Variant Calling Format
*CHH*: Congenital hypogonadotropic hypogonadism
*CPHD*: Combined pituitary hormone deficiency
*CHARGE syndrome*: Coloboma, heart defects, atresia choanae, growth retardation, genital abnormalities, and ear abnormalities.

## Ethics approval and consent to participate

Not applicable

### Consent for publication

Not applicable

## Availability of data and material

GenMasterTable is available under the MIT license at https://github.com/strawberrybeijing/GenMasterTable. The repository also includes the full source code of the application along with artificially generated test datasets (VCF and CSV files) used for the demonstration in the manuscript.

## Authors’ contributions

NP conceptualized the study, contributed to the overall design, and supervised the development and evaluation of the software. FS provided expert guidance on the study design, data processing strategies, and interpretation of genomic datasets. JZ developed and implemented the GenMasterTable software. All authors read and approved the final manuscript.

## Acknowledgements

Not applicable

## Authors’ information

Faculty of Biology and Medicine, University of Lausanne, Lausanne, Switzerland Jing Zhai, Nelly Pitteloud, Federico A. Santoni

Service of Endocrinology, Diabetology and Metabolism, Lausanne University Hospital, Lausanne, Switzerland Jing Zhai, Nelly Pitteloud, Federico A. Santoni

Federico A. Santoni and Jing Zhai are co-corresponding authors.

## References

1. McLaren W, et al. The Ensembl Variant Effect Predictor. Genome Biol. 2016;17:1–14.

2. Wang K, et al. ANNOVAR: functional annotation of genetic variants from high-throughput sequencing data. Nucleic Acids Res. 2010;38:e164.

3. Zhou H, et al. FAVOR: functional annotation of variants online resource and annotator for variation across the human genome. Nucleic Acids Res. 2023;51:D1300–11.

4. Paila U, et al. GEMINI: integrative exploration of genetic variation and genome annotations. PLoS Comput Biol. 2013;9:e1003153.

5. Wang GT, et al. Variant association tools for quality control and analysis of large-scale sequence and genotyping array data. Am J Hum Genet. 2014;94:770–83.

6. Cingolani P, et al. Using Drosophila melanogaster as a model for genotoxic chemical mutational studies with a new program, SnpSift. Front Genet. 2012;3:35.

7. Pietrelli A, Valenti L. myVCF: a desktop application for high-throughput mutations data management. Bioinformatics. 2017;33:3676–8.

8. Salatino S, Ramraj V. BrowseVCF: a web-based application and workflow to quickly prioritize disease-causative variants in VCF files. Brief Bioinform. 2017;18:774–9.

9. Hart SN, et al. VCF-Miner: GUI-based application for mining variants and annotations stored in VCF files. Brief Bioinform. 2016;17:346–51.

10. Akgun M, Demirci H. VCF-Explorer: filtering and analysing whole genome VCF files. Bioinformatics. 2017;33:3468–70.

11. Jiang J, et al. VCF-Server: a web-based visualization tool for high-throughput variant data mining and management. Mol Genet Genomic Med. 2019;7:e00641.

12. Schutz S, et al. Cutevariant: a standalone GUI-based desktop application to explore genetic variations from an annotated VCF file. Bioinform Adv. 2021;1:vbab028.

13. Zaharia M, et al. Spark: cluster computing with working sets. Proc 2nd USENIX Conf Hot Topics Cloud Comput (HotCloud’10). 2010.

14. Young J, et al. Clinical management of congenital hypogonadotropic hypogonadism. Endocr Rev. 2019;40:669–710.

