## Supplemental File 1 for "GenMasterTable: A user-friendly desktop application for filtering, summarising, and visualising large-scale annotated genetic variants"

### GenMasterTable Handbook

Jing Zhai

7th March 2025

### Introduction

GenMasterTable is a user-friendly graphical user interface (GUI) designed to facilitate the visualization, filtering, and summarization of **large-scale datasets** that are difficult to manage with conventional spreadsheet applications like Excel. A key motivation behind GenMasterTable is to bridge the gap for non-technical users, such as clinicians, geneticists, and researchers, who may not have experience with databases (e.g., Oracle, MSSQL, PostgreSQL) or programming languages (e.g., Python, R) commonly used for large-scale data analysis. By offering an intuitive GUI, GenMasterTable enables efficient data exploration, filtering, and transformation without requiring expertise in SQL queries or scripting with data-processing libraries such as pandas.

This handbook provides a comprehensive guide on how to use GenMasterTable, covering its core functionalities with step-by-step instructions. To demonstrate its capabilities, we include a demo dataset comprising artificially generated genomic data in VCF and CSV formats, allowing users to test features such as merging, filtering, summarising, and visualization in genomic date. These demo datasets can be found in the GenMasterTable GitHub repository.

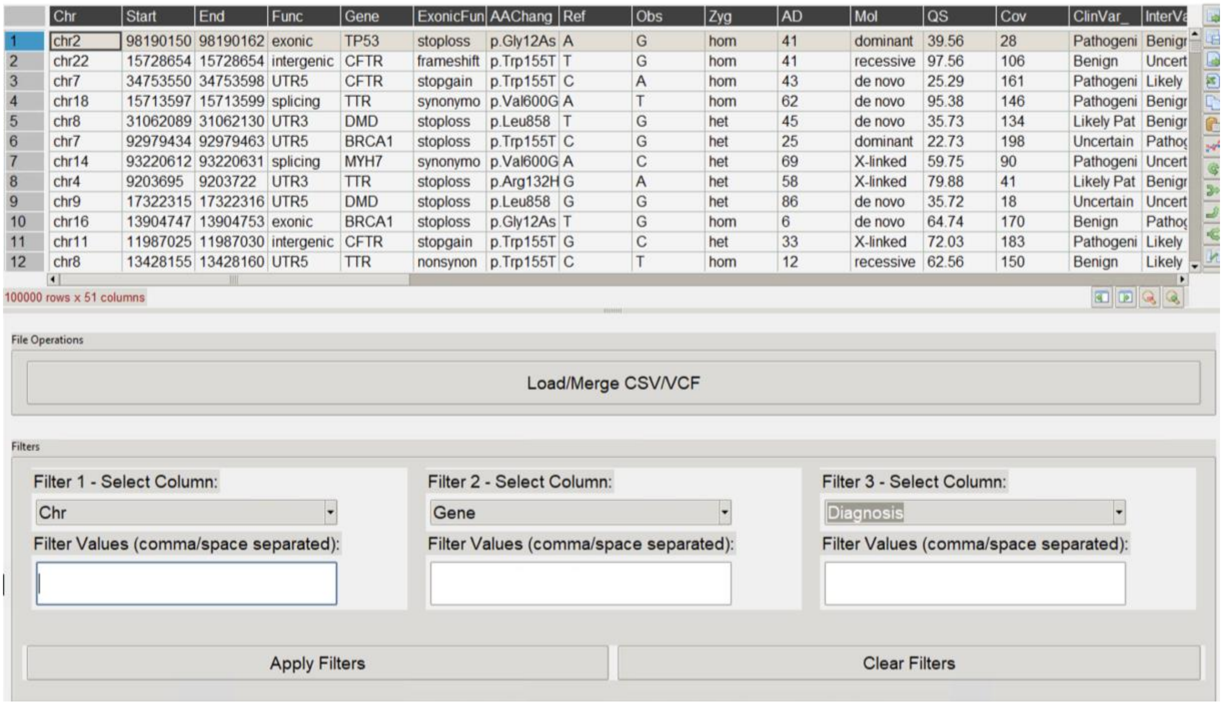

#### 1.1Installation

To install GenMasterTable, visit the GitHub **Releases** page and download the latest version compatible with your operating system:

- Windows 64-bit: Download **GenMasterTable_windows.exe**
- macOS: Download **GenMasterTable_MacOS.zip**

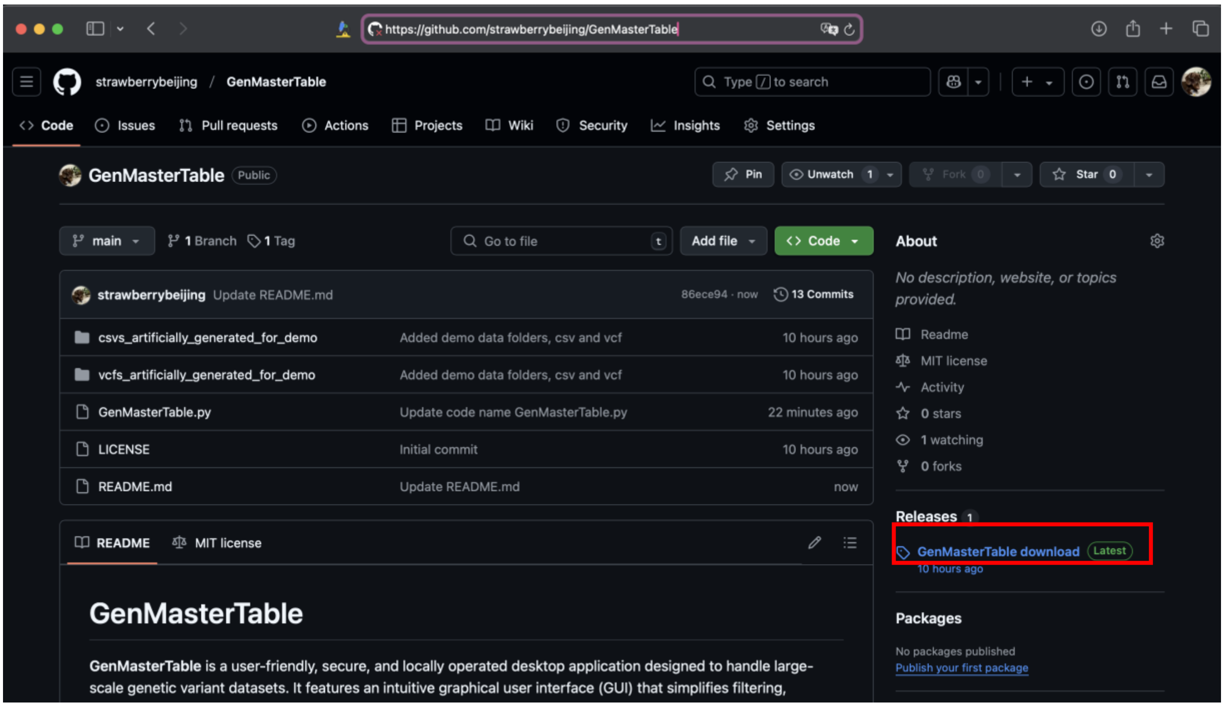


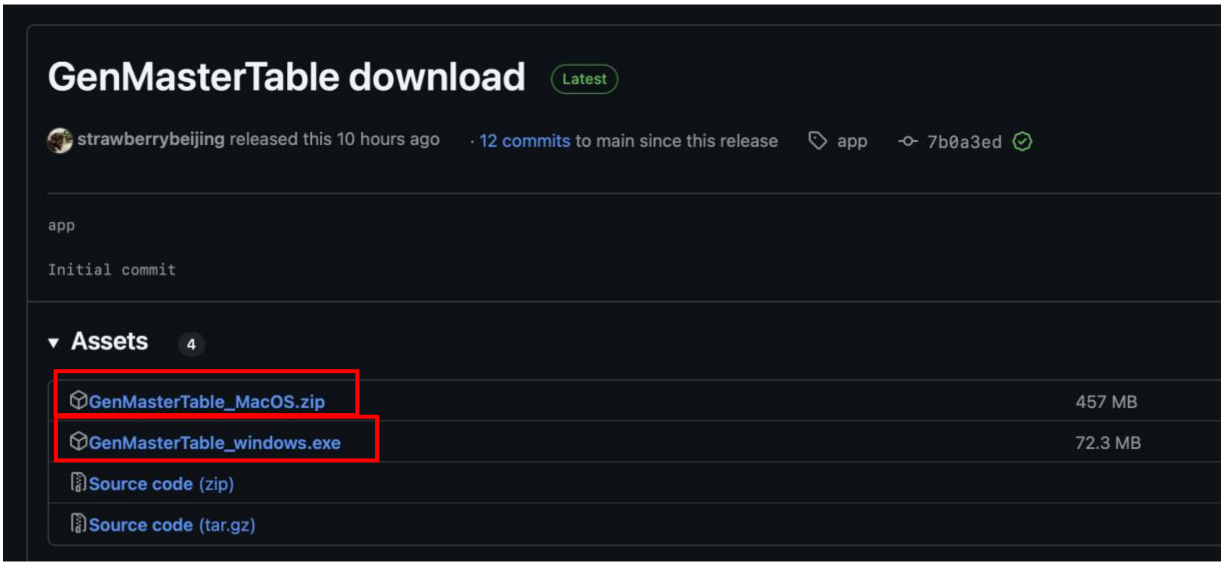

#### 1.2 Launching GenMasterTable

Double-click the downloaded GenMasterTable, just as you would to open any other software. If you encounter a security warning, navigate to System Preferences > Security & Privacy and allow the app to run.

### 2. Load and Merge tables

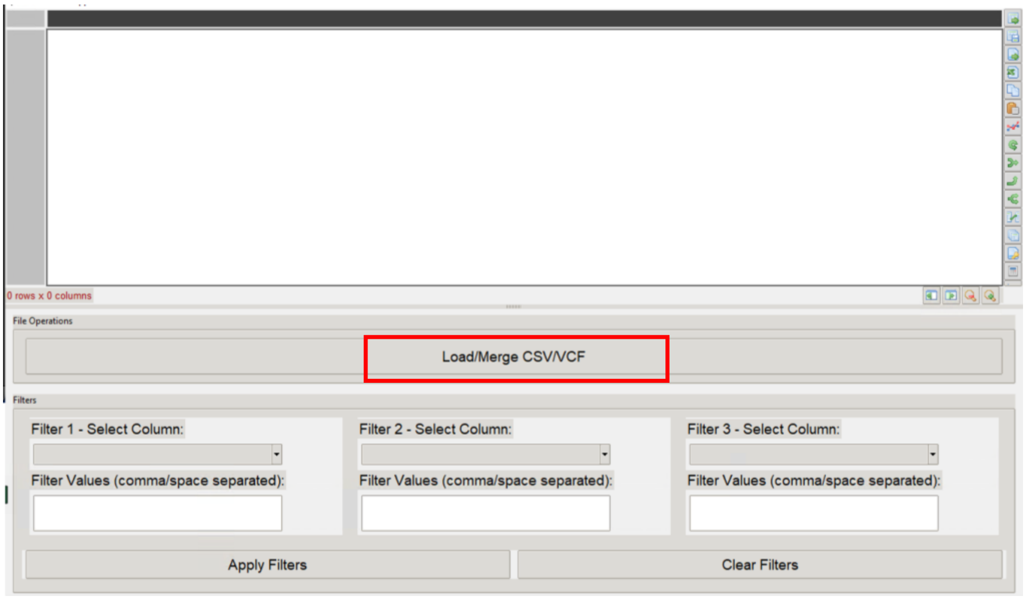


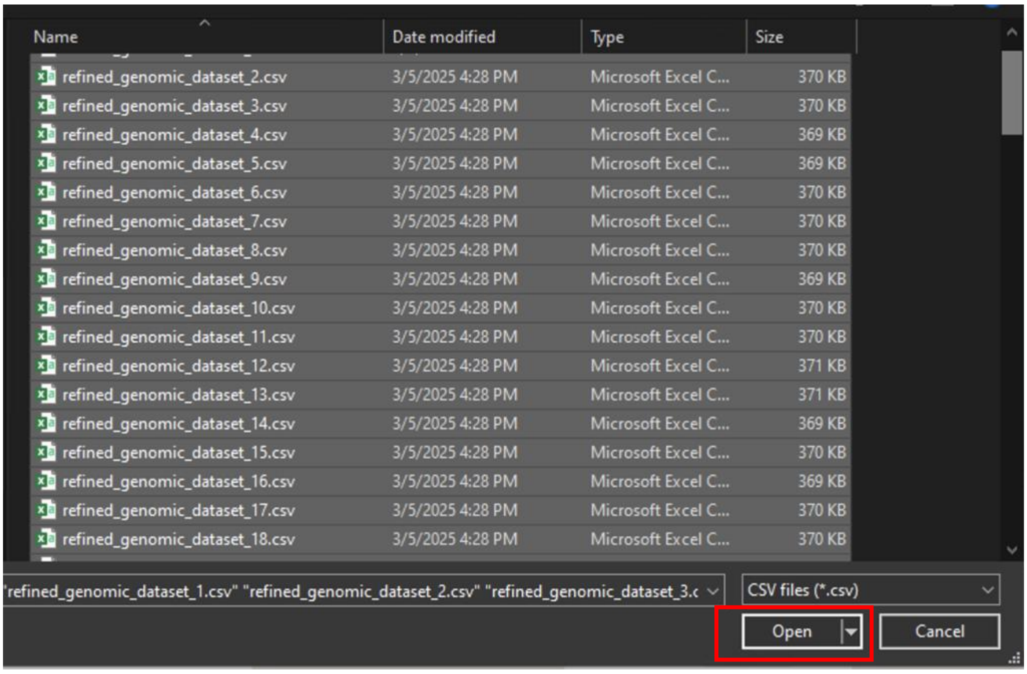


Once the software is launched, click the central button to select the dataset you want to work with. You can select a single VCF/CSV file or choose multiple files to merge into a larger dataset for analysis. In this demo, I will select 100 artificially generated CSV files (available in the GitHub repository) to merge and load for demo.

### 3. Filter functions

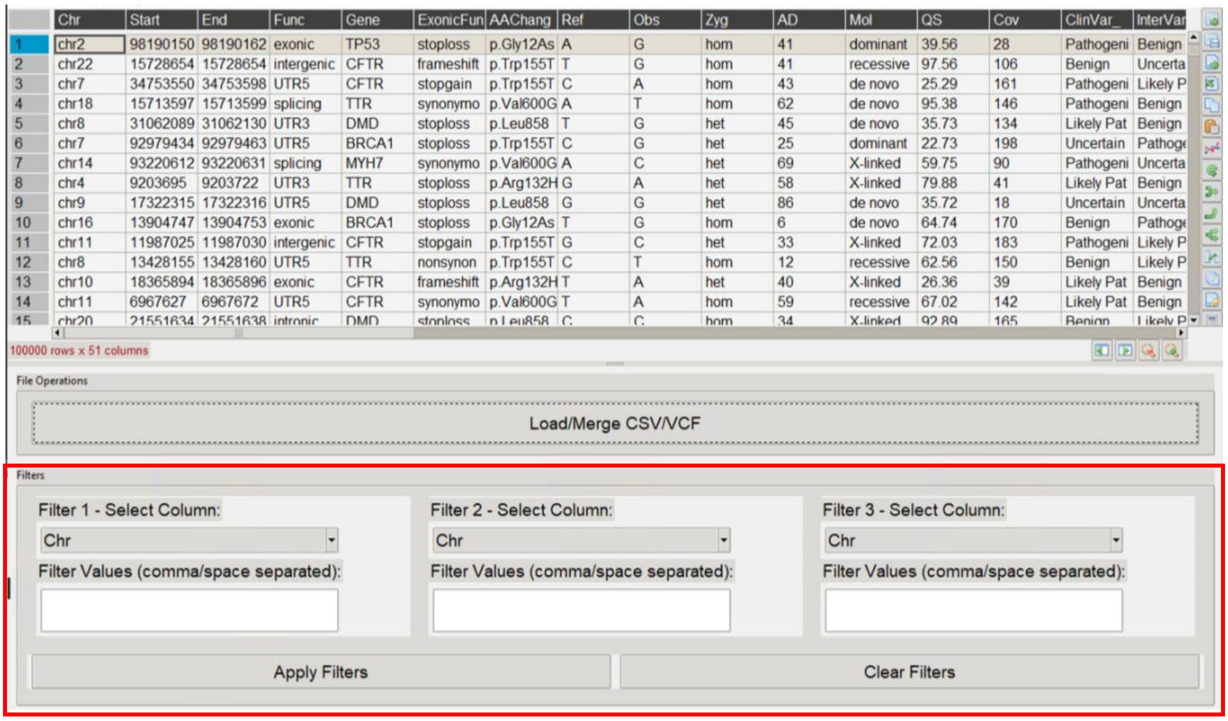


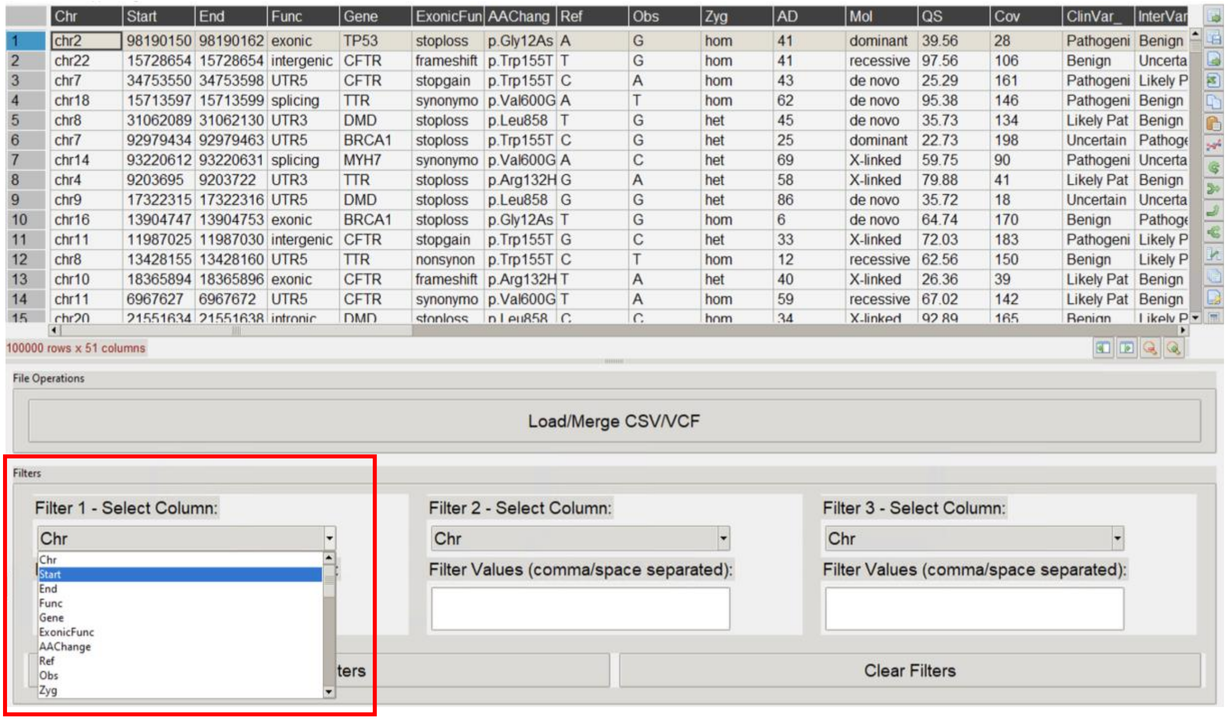

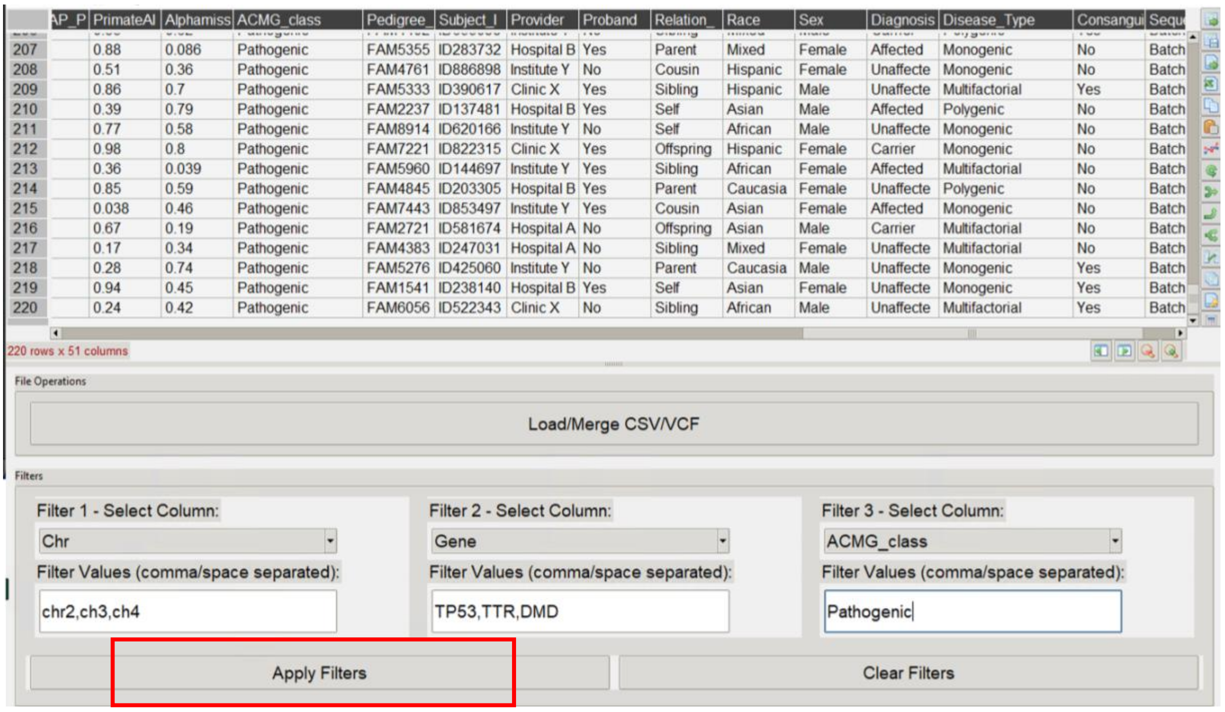


The main filtering panel, located in the control frame, displays a scrollable list of columns, allowing you to select the target columns for filtering. Filter criteria should be separated by a comma or space when specifying multiple values. Once all selections are made, click "**Apply Filters**" to apply them. To remove all filters, click "**Clear Filters**".

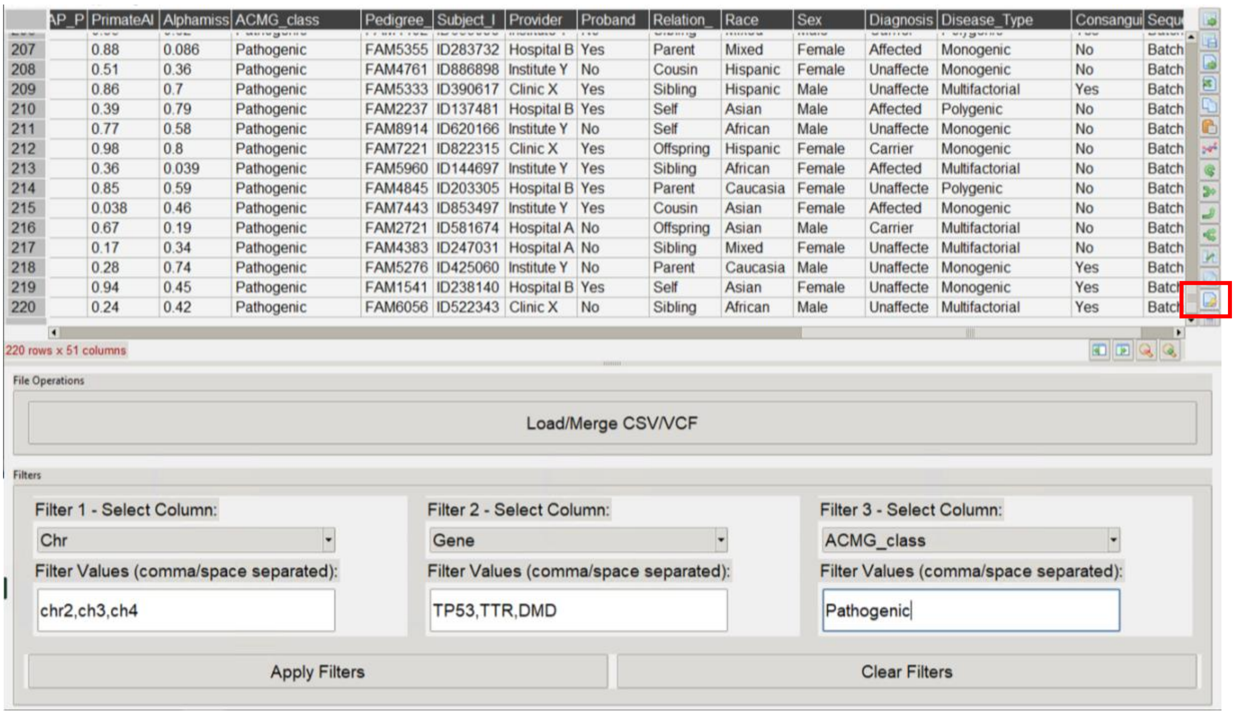


Furthermore, GenMasterTable allows filtering based on quantitative thresholds using the additional filter function located on the right side of the interface. Click the filter button with a key icon, enter the target quantitative column, and define the threshold criteria. When applying filters across multiple columns, conditions can typically be set using AND or OR operators.

Difference between **AND & OR** among multiple filters/conditions:

- The **AND** operator is used to combine multiple conditions in a way that all of them must be true for a row to be included in the query result.
- The **OR** operator is used to combine multiple conditions in a way that at least one of them must be true for a row to be included in the query result.

Click the "Apply Filters" button located just below the filter criteria to execute the filtering process:

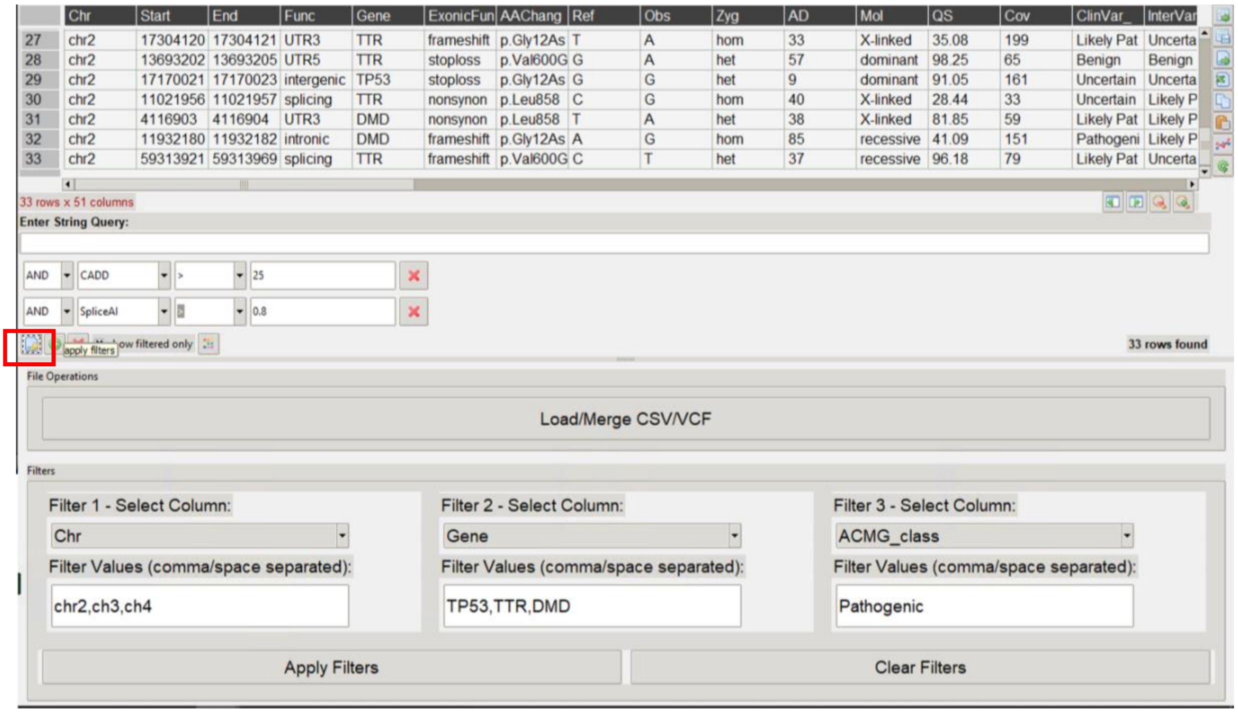


**Troubleshooting Filter Issues with Quantitative Variables:**

If you encounter difficulties with filters not functioning as expected for a quantitative variable, it's important to ensure that the column is populated with numerical data, specifically using the data types 'float64' (which correspond to numerical data in human terms), rather than 'str' or 'object' (indicating textual data).

To rectify this, you can follow these steps:

- Right-click on the column in question.
- Locate the 'Set Data Type' option and select it.
- Choose 'float64' from the available options to set the correct data type for the column.

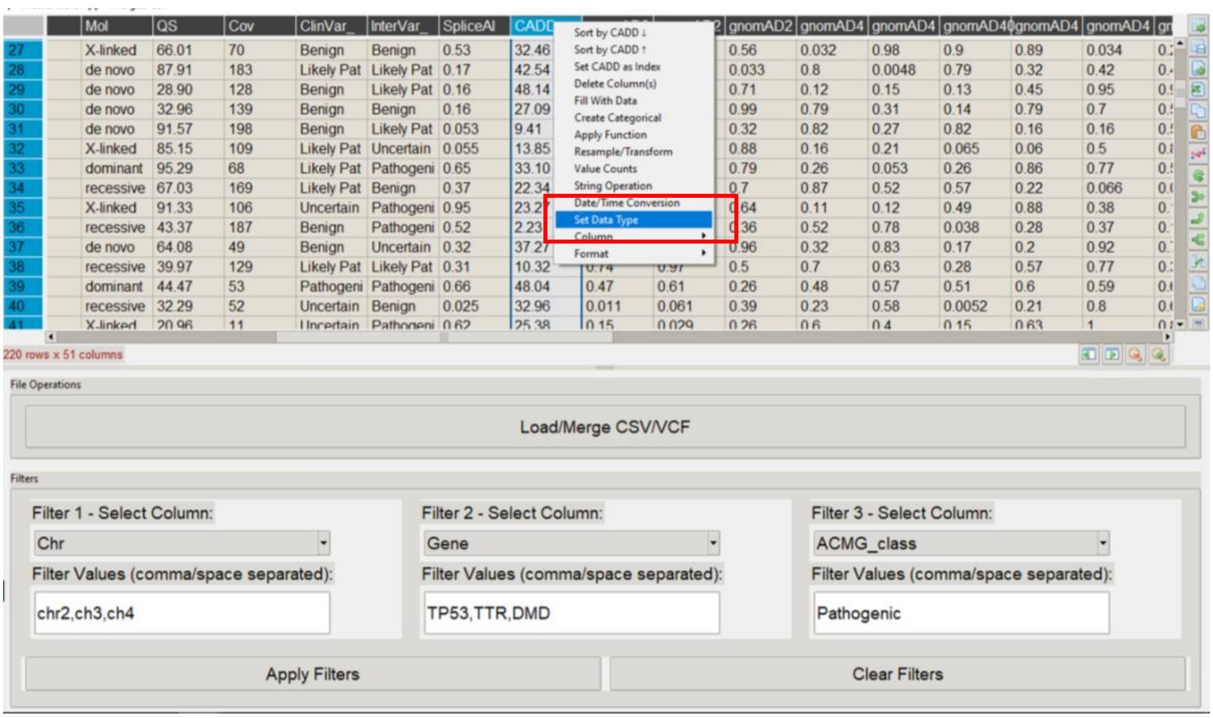
Sort by CADD ↓
Sort by CADD ↑
Set CADD as Index
Delete Column(s)
Fill With Data
Create Categorical
Apply Function
Resample/Transform
Value Counts
String Operation
Date/Time Conversion
Set Data Type
Column
Format
220 rows x 51 columns
File Operations
Load/Merge CSV/VCF
Filters
Filter 1 - Select Column:
Chr
Filter Values (comma/space separated):
chr2,ch3,ch4
Filter 2 - Select Column:
Gene
Filter Values (comma/space separated):
TP53,TTR,DMD
Filter 3 - Select Column:
ACMG_class
Filter Values (comma/space separated):
Pathogenic
Apply Filters
Clear Filters

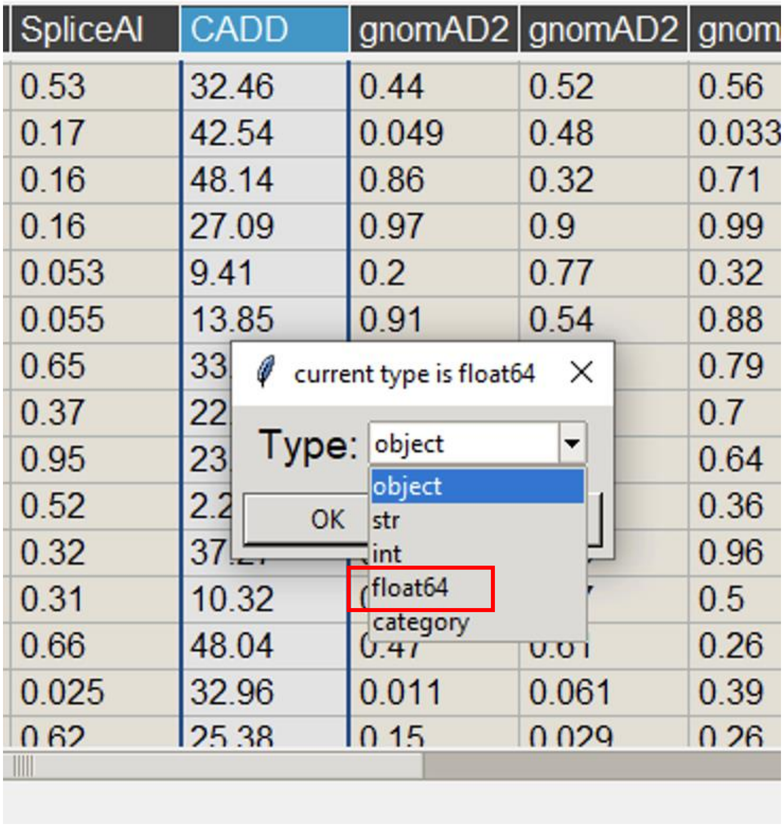
SpliceAI
CADD
gnomAD2
current type is float64
Type:
object
str
int
float64
category
OK

#### 4. Data Export

After filtering subsets or merging CSV and VCF files, GenMasterTable allows users to copy and paste the data directly into an open Excel sheet. First, click the top-left corner of the working sheet to select all data.

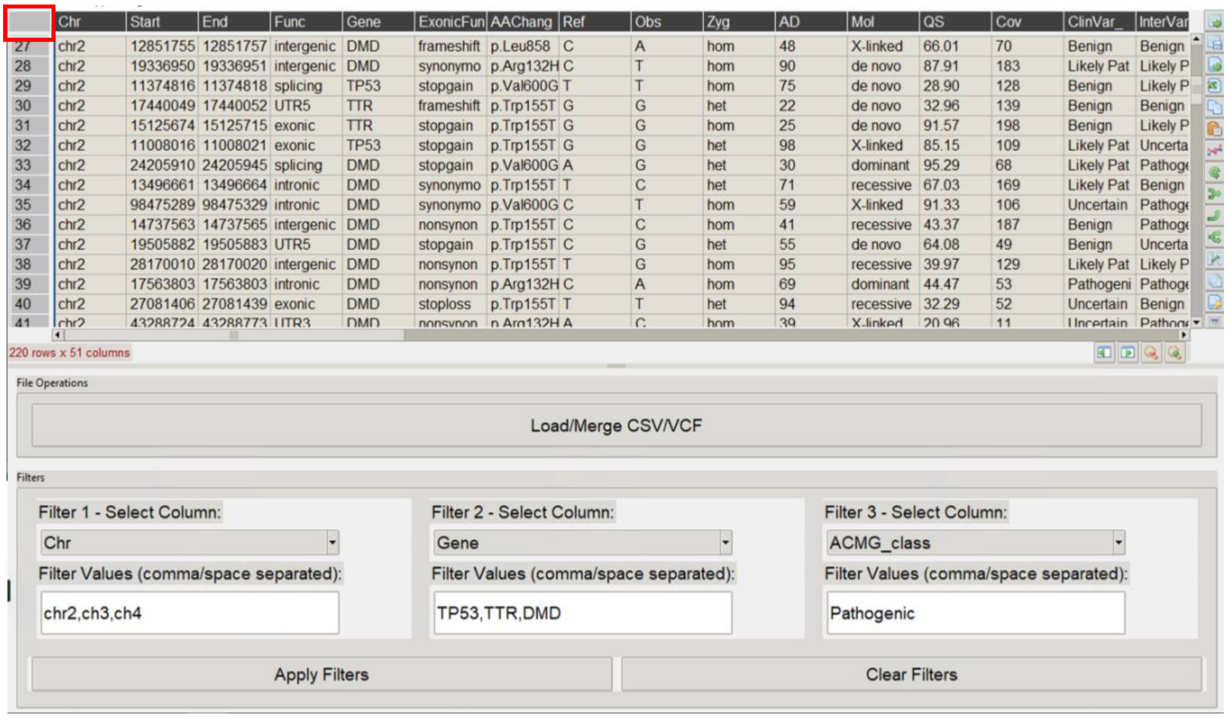


Then, click the copy icon (on the right side of the interface) to copy the selected data. The data is now ready to be pasted directly into an open Excel sheet.

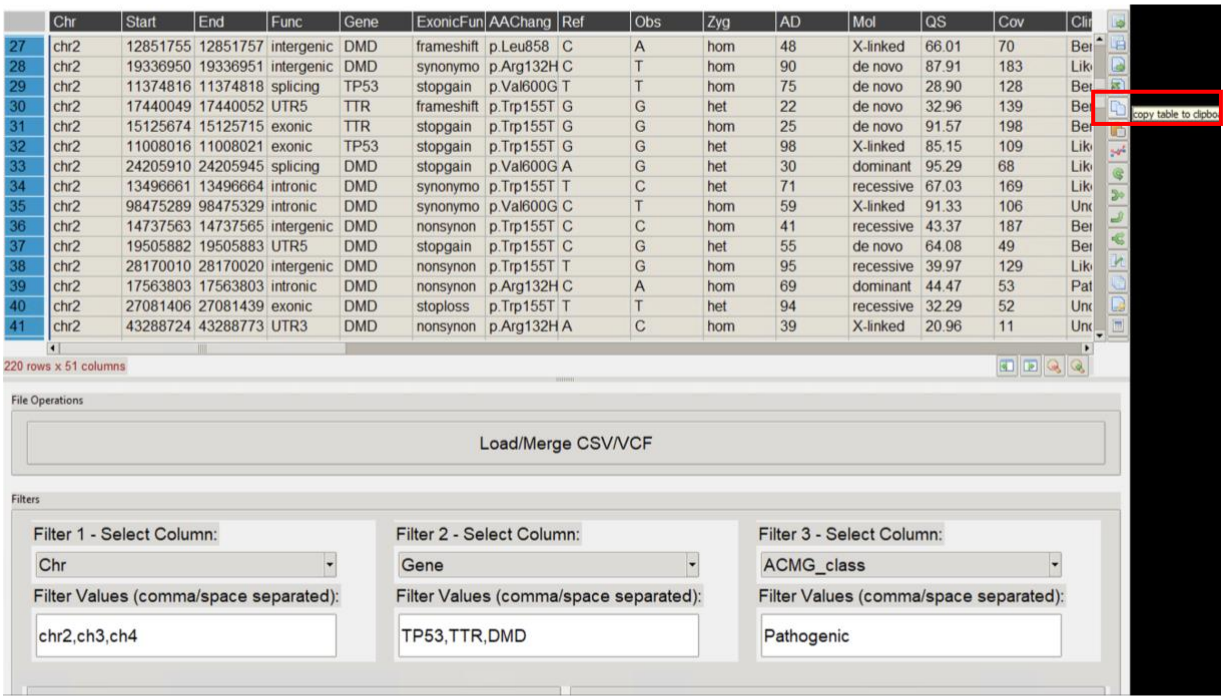

#### 5. Additional Functions for Using the GenMasterTable

To enhance table visualization, users can use the "Zoom In" and "Zoom Out" functions to adjust the display for better readability and analysis.

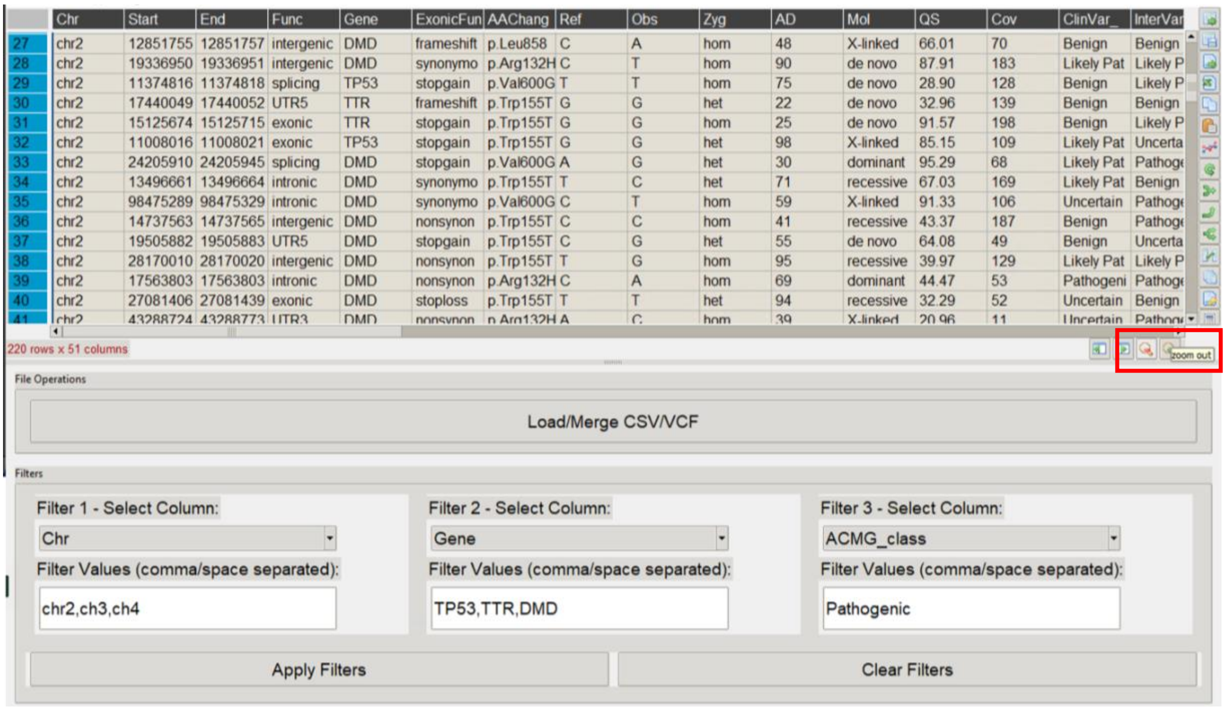

By right-clicking on a column header, users can access additional data visualization and manipulation options. For instance, columns can be sorted in ascending or descending order, enabling efficient data organization.

For a practical demonstration, let's explore how to identify variants with the **highest** AlphaMissense pathogenicity scores within our cohort.

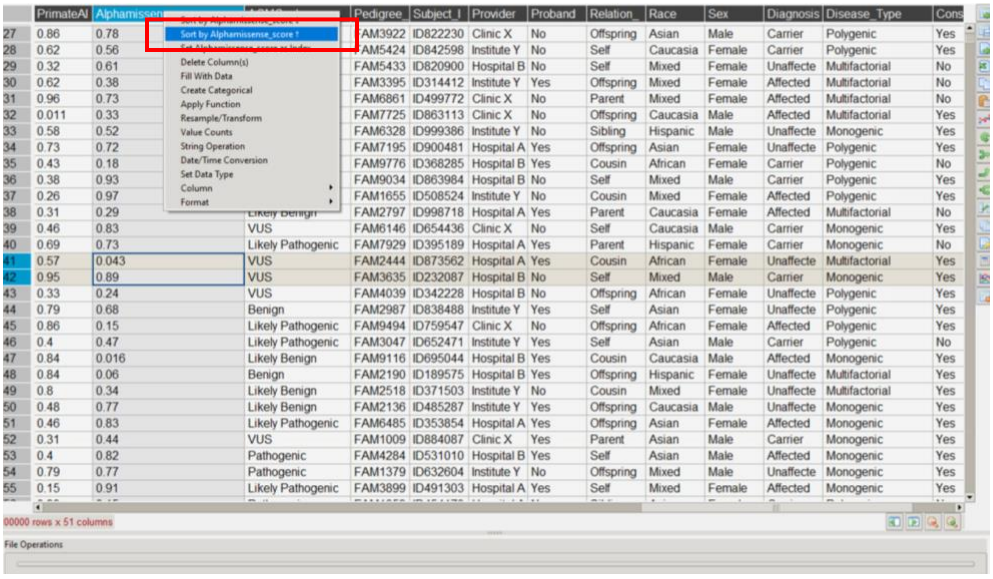


Users can also hide unnecessary columns by using the "Delete Columns" function, allowing for a more focused analysis. It is important to note that deleting columns does not affect the original dataset—this action only applies to the current working session, as GenMasterTable operates as a GUI-based tool without modifying the source data.

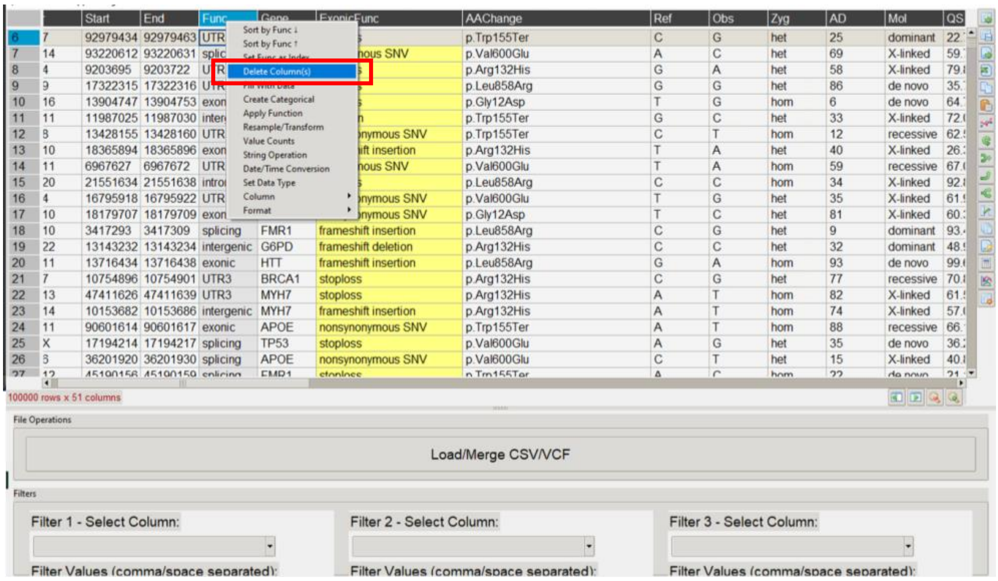

Another valuable feature available to users is the "Value Counts" function, which provides a frequency distribution of data within a selected column.

For example, we can analyze the number of variants per subject ID within our cohort. In this case, the dataset shows that we have 94,786 subjects, with most individuals carrying at most four variants.

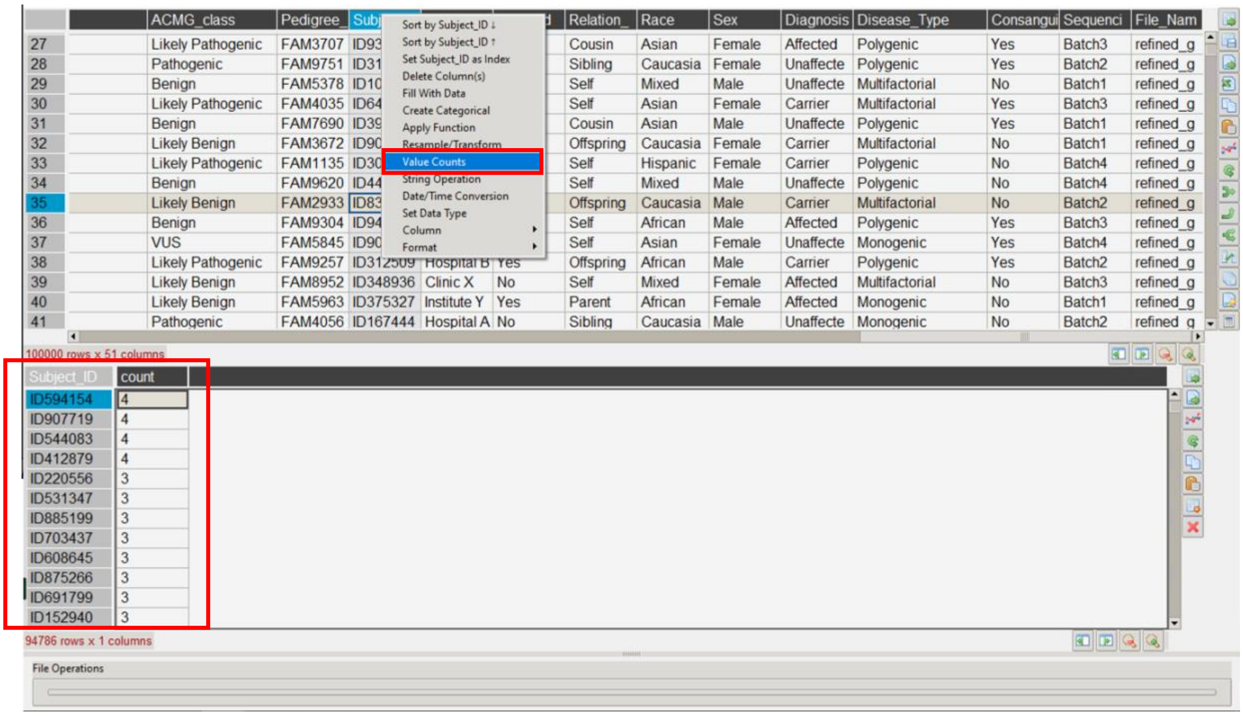


GenMasterTable also provides a column highlighting feature, allowing users to apply color applies to specific columns for improved data visualization and readability.

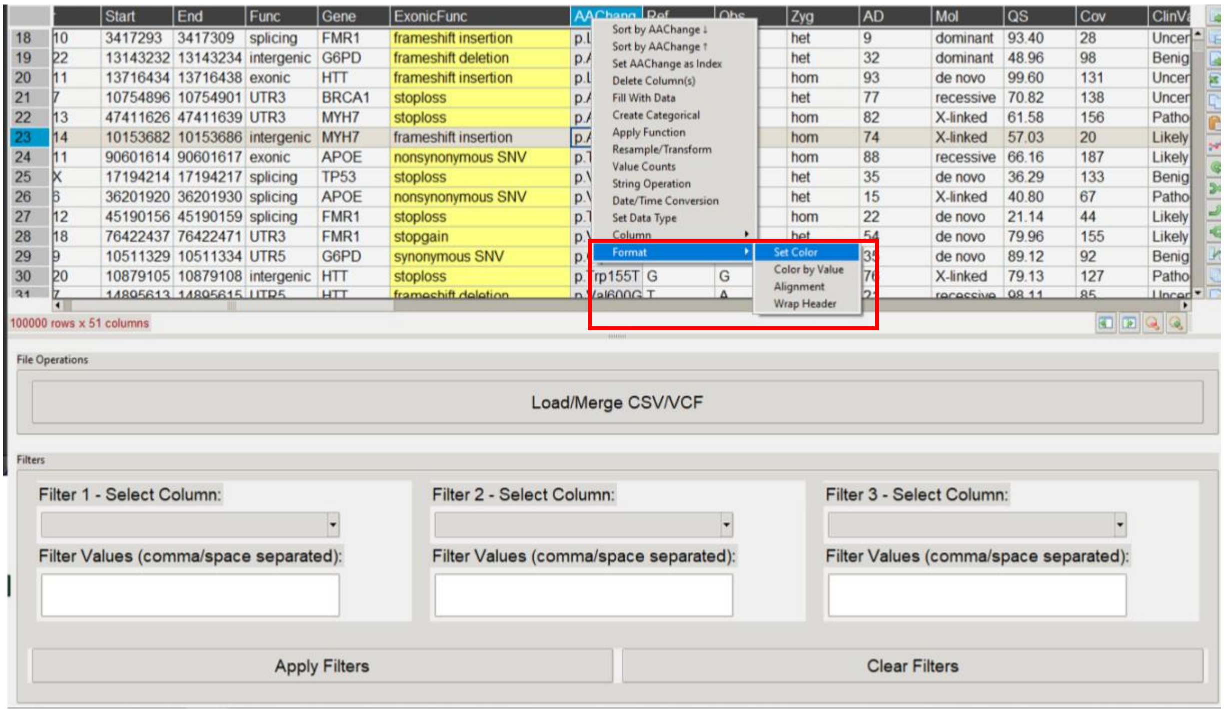

The order of columns can be adjusted, allowing users to move any column to the first or last position. Additionally, column names can be renamed for better clarity and organization.

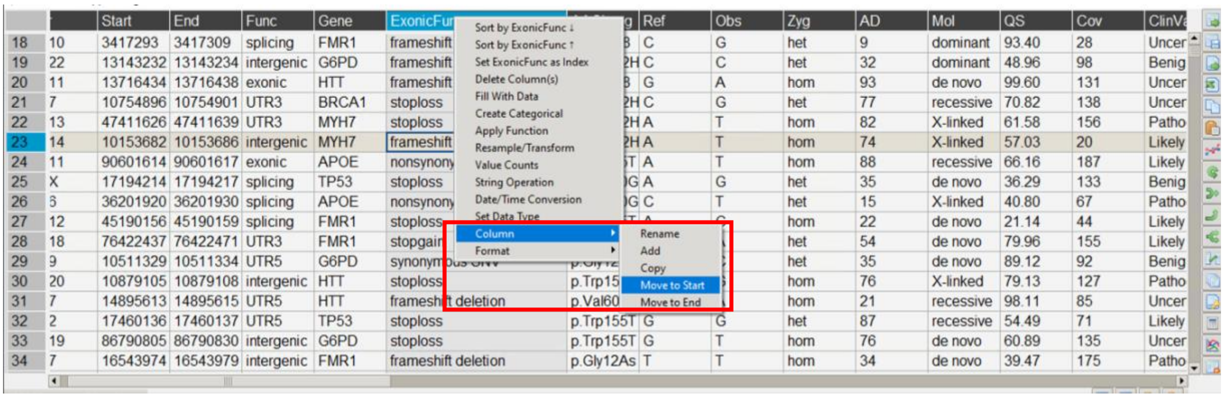


#### 6.Conclusion:

Throughout this handbook, we have demonstrated how to install, load, merge, filter, and export data using GenMasterTable, along with additional features such as value counts, quantitative filtering, column manipulation, and data visualization enhancements. These functionalities enable users to navigate and analyze large datasets with ease, ensuring efficient cohort-level analyses.
